# Context-aware simulation enables systematic optimization of long-read mapping parameters

**DOI:** 10.64898/2025.12.04.692264

**Authors:** Jiang Hu, Dongming Fang, Xin Jin, Chentao Yang

**Affiliations:** BGI Research, Wuhan 430074, China; State Key Laboratory of Genome and Multi-omics Technologies, BGI Research, Shenzhen 518083, China; Center for Evolutionary Biology, School of Life Sciences, Fudan University, Shanghai 200438, China; Guangdong Provincial Key Laboratory of Genome Read and Write, BGI Research, Shenzhen 518083, China

**Keywords:** Long-read sequencing, long-read simulator, Context-aware simulation, Parameter tuning, Bayesian optimization

## Abstract

The performance of long-read mapping is critical yet highly sensitive to parameter choices. We present CycSim, a context-aware simulator that models sequence-context-dependent errors from empirical sequencing data, coupled with a Bayesian optimization framework for systematic parameter tuning. CycSim more accurately reproduces real error profiles than existing simulators, enabling reliable simulation-based optimization. The framework identified parameter configurations that achieved 2.78-fold faster mapping for data from the newly developed Cyclone platform, and consistently improved both mapping efficiency (8.14-32.65% faster) and structural variant calling accuracy (0.75-1.70% higher F1) across ONT, HiFi, and Cyclone datasets, providing a robust and generalizable foundation for analysis-goal-driven parameter refinement.

## Background

The accuracy of long-read sequence mapping is fundamental to genomic analysis but is sensitive to parameterization [1–3]. Default parameters offer convenience but are often suboptimal, failing to generalize across sequencing platforms such as PacBio high-fidelity [4] (HiFi), Oxford Nanopore Technologies [5] (ONT), and Cyclone [6], or across analytical objectives such as structural variant (SV) detection. Systematic optimization is impeded by the empirical nature of default parameters, the scarcity of datasets with known ground-truth alignments, and the lack of a comprehensive framework for identifying optimal settings for specific data types or analyses. Simulation-based evaluation offers a practical solution, yet existing long-read simulators largely introduce errors at random, failing to capture sequence-context-dependent biases and global error rate heterogeneity characteristic of real data [7–9]. As a result, current simulators provide limited realism for benchmarking and parameter tuning.

To address this, we developed CycSim, a context-aware long-read simulator that generates reads based on K-mer contexts and error distributions learned from empirical datasets. We integrated CycSim with a Bayesian optimization framework to systematically identify optimal mapping parameters. Benchmarking confirms that CycSim reconstructs error profiles with greater realism than state-of-the-art alternatives. Furthermore, our optimization framework identified superior parameter sets for the Cyclone platform and confirmed that the robustness of default settings for HiFi and ONT data. Notably, SV-specific tuning improves both alignment speeds and accuracy (F1 scores) across all three platforms.

## Results and Discussion

### Context-aware long-read simulation

CycSim operates through a dual-stage framwork comprising model training and read simulation (**Fig. S1**). During the training stage, the algorithm charaterizes read structure, including strand orientation, chimerism, aligned/unaligned lengths, and alignment identity by interrogating high-confidence alignment from input BAM files. To mitigate artifacts arising from alignment heuristics, aligned regions are re-aligned using edlib [10] and low-confidence termini are trimmed. Crucially, we model error characteristics at two complementary levels: (1) K-mer-based error modeling, which captures the empirical frequency of context-dependent substitution, insertion, and deletion via a sliding window approach; and (2) error transition modeling, which estimates the transition probabilities between consecutive error states to reflect the local continuity of error types.

In the simulation stage, these models define the genomic origin, structural composition, and expected error rate of synthetic reads. The aligned core is generated through a base-wise sliding process, in which errors are sampled according to the K-mer specific models and the empirical error transition matrix. Following this, Phred-scaled quality scores are then assigned, and unaligned regions are appended to form complete reads. When the simulated error rate deviates markedly from the expected value, resampling is performed, and chimeric reads are constructed by concatenating independent simulated fragments.

To validate the framework, we benchmarked CycSim against BadRead [9], NanoSim [7], and PbSim3 [8] using the diploid HG002 genome [11] (Chromosomes 1, 2, 17, and 18) accross ONT, HiFi and Cyclone platforms. CycSim consistently demonstrated superior fidelity in reproducing both read length and global error rate distributions (**Fig. 1A**, **Fig. S2**). Our assessment focused on three key metrics: (1) Base substitution profiles: CycSim and BadRead were the only tools to accurately capture empirically observed substitution biases (**Fig. 1B**); (2) K-mer distribution: CycSim faithfully reproduced the global K-mer frequency landscape. specifically, it achieved the highest concordance for erroneous K-mers in Cyclone and HiFi data, while performing competitively on ONT (**Fig. 1C**); (3) Error rates in simple repeats: Notably, real Cyclone and ONT data exhibit substantial error rate heterogeneity in low-complexity regions. While other simulators produced artifactually uniform error distributions. CycSim successfully recapitulated these localized, context-dependent error trends (**Fig. 1D, Fig. S3**).

**Figure 1.**
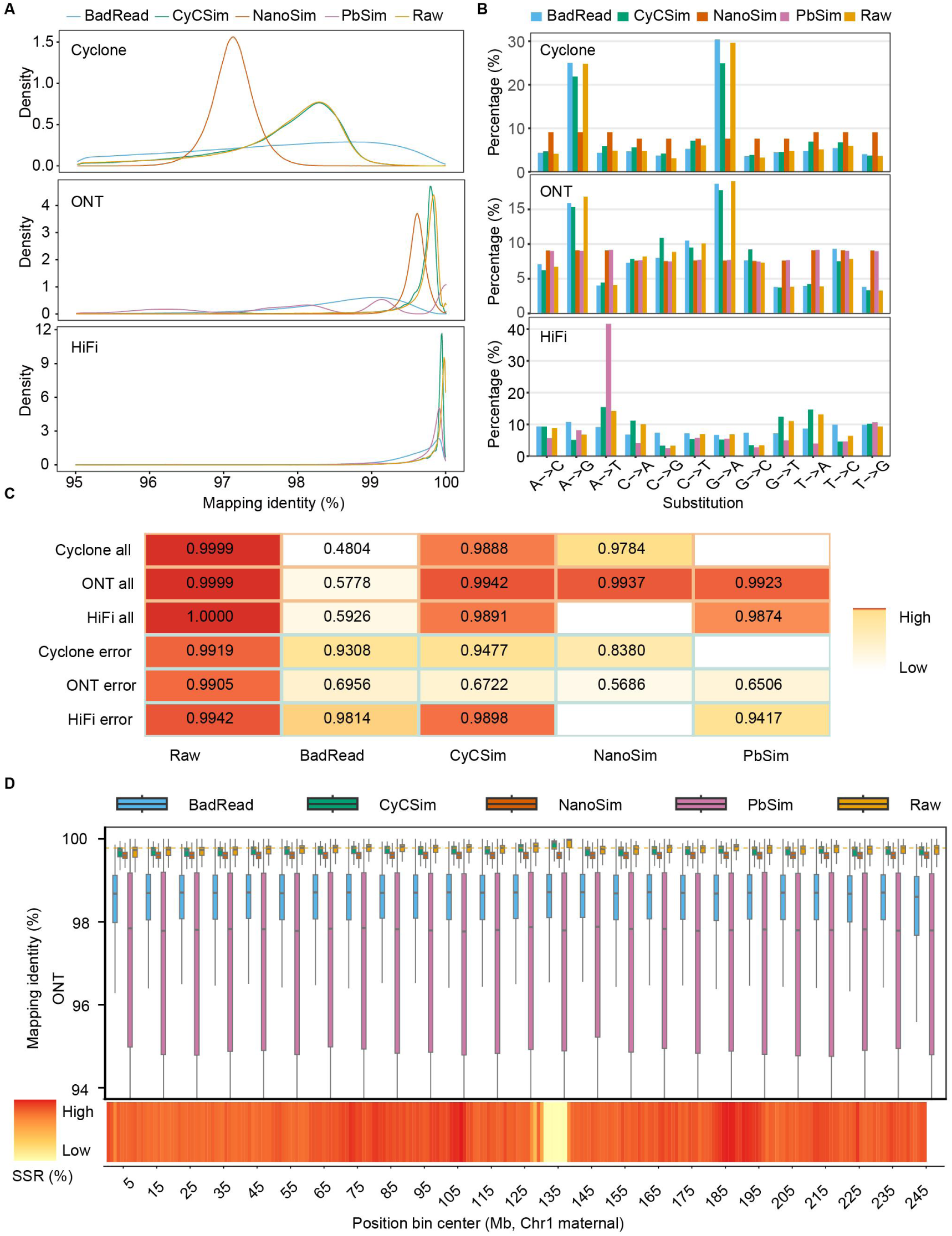
Multidimensional evaluation of long-read simulators. **A** Distribution of alignment identity for simulated reads compared with real reads (Raw). **B** Statistics of substitution error bias of long-read simulators. **C** Cosine similarity of K-mer count between simulated and real reads. All denotes similarity computed over all K-mers; Error denotes K-mers present in the real reads but absent from the reference genome, representing sequencing-induced errors. For Raw, real reads were randomly split into two parts to estimate an empirical upper bound of similarity. Blank entries indicate missing values. **D** Positional alignment identity distribution along Chr1 maternal for simulated ONT reads and real reads. Horizontal yellow lines mark the median identity of real ONT reads. The lower heatmap shows the short tandem repeats (STRs) density (1–6 bp motifs, ≥3 repeat units) along Chr1 maternal.

Collectively, these analyses demonstrate that CycSim provides a high-fidelity, platform-consistent representation of long-read error characteristics, capturing both global and localized, context-dependent biases that existing simulators fail to model.

### Bayesian optimization of mapping parameters

Leveraging the fidelity of CycSim-generated data, we established a four-stage Bayesian optimization framework to systematically identify optimal mapping parameters. The workflow proceeds as follows: (1) Simulation-based initialization using CycSim-generated reads with known ground-truth coordinates; (2) Bayesian parameter search via Optuna [12], which explore thousands of minimap2 [2] configurations to maximize a defined objective function; (3) Empirical screening, where top-performing parameter sets are evaluated on real data subsets. (4) Whole-genome validation to ensure robustness across full-scale datasets.

We first applied the framework to optimize general-purpose alignment (**Fig. 2A**). The optimization utilized 5× CycSim-simulated HG002 data (Chr 1, 2, 17, 18) to maximize a composite accuracy metric integrating both base-level identity and interval-level overlap. From this search, the top 40 configurations were screened using 30× real HG002 data on the same chromosomal subset, assessed using standard SNP and SV benchmarks. The optimal configuration was subsequently validated on 44× whole-genome data.

**Figure 2.**
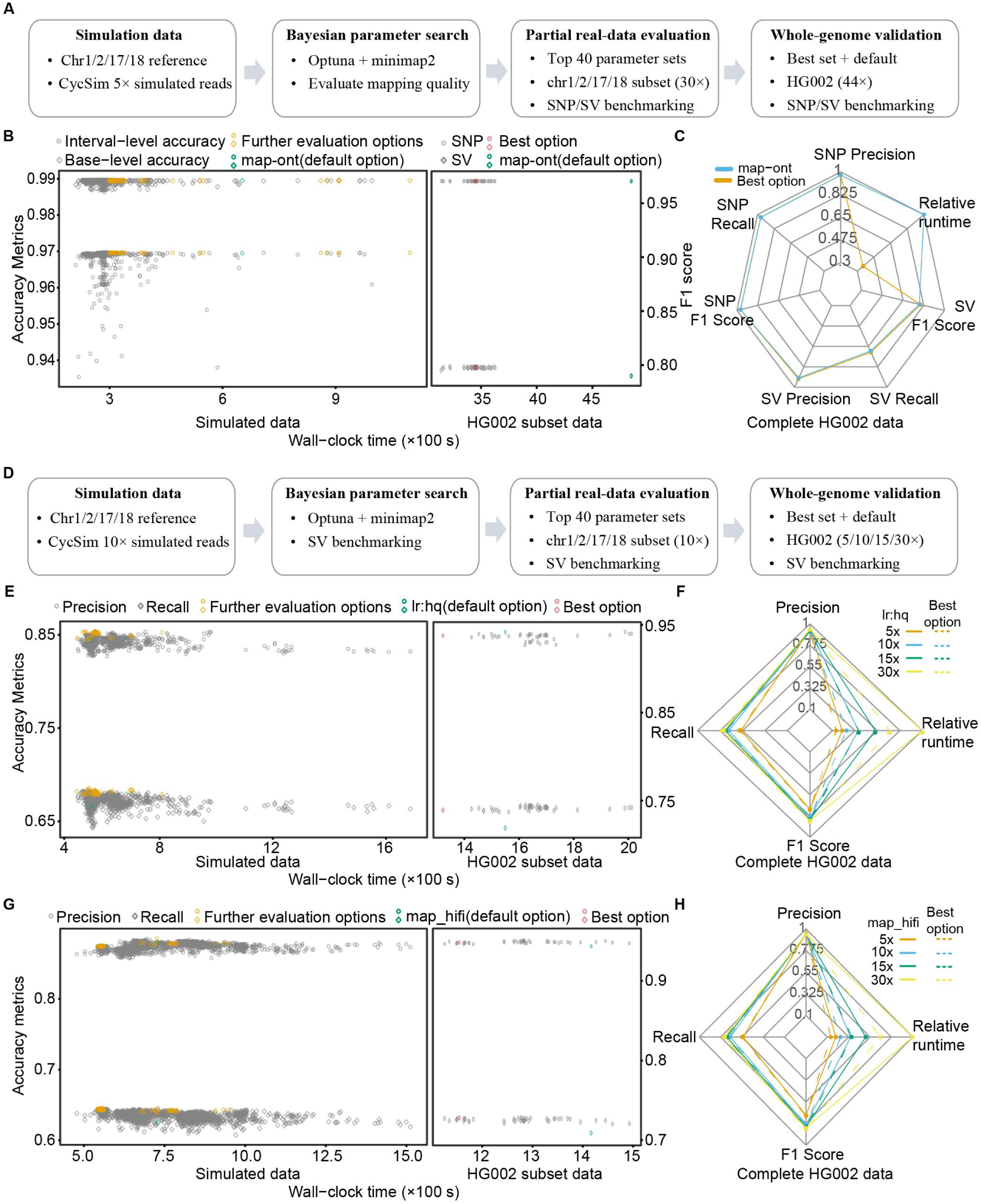
Bayesian optimization and evaluation of mapping parameters. **A** Optimization workflow for general-purpose mapping parameters. **B** Performance of Cyclone mapping parameters on simulated and partial real reads. Interval-level accuracy denotes the proportion of aligned intervals within 50 bp of the true interval, and Base-level accuracy denotes the proportion of correctly aligned bases. SNP and SV accuracy were assessed as described in Methods. **C** Performance of optimized versus default parameters on 44× real Cyclone reads. **D** Optimization workflow for SV detection-oriented mapping parameters. **E** Performance of ONT SV detection-oriented mapping parameters on simulated and partial real reads. **F** Performance of optimized versus default ONT mapping parameters for SV detection at different coverage depths. **G** Performance of HiFi SV detection-oriented mapping parameters on simulated and partial real reads. **H** Performance of optimized versus default HiFi mapping parameters for SV detection at different coverage depths. For Panels B, E, and G, parameters evaluated in the right panel were selected from the left panel (labeled as Further evaluation options), and Wall-clock time includes both minimap2 and samtools sort. Relative runtime (Panels C, F, H) denotes the proportion of runtime relative to the longest run, considering minimap2 only.

When applied to the emerging Cyclone platform (using the map-ont preset as a baseline), our framework identified a novel parameter set that increased mapping speed by 2.78-fold while maintaining comparable SNP-calling accuracy and modestly enhancing SV detection (**Figs. 2B and 2C**). Conversely, for HiFi and ONT datasets, the framework confirmed that default presets are near-optimal, our best-performing configurations yielded marginal accuracy gains (<0.1% in simulation) with slightly reduced runtime (**Fig. S4**). These results demonstrate the framework’s robustness in reliably optimizing parameters across diverse sequencing platforms or emerging chemistries.

We next adapted the framework specifically for SV detection, a task strictly reliant on interval-level alignment accuracy (**Fig. 2D**). While retaining the four-stage process, we modified the objective function to maximize the SV F1-score. Top candidate parameters were subsequently screened and evaluated using SV benchmarks on both chromosomal subsets and whole-genome datasets.

Across all three platforms, SV-specific optimization yielded configurations efficiency compared to default settings. Notably, on whole-genome HG002 datasets at varying depths, the optimized parameters achieved significant performance gains: 30.84-32.65% faster mapping and 0.75-1.37% higher SV F1-scores for ONT data (**Figs. 2E and 2F, Table S1**), 28.34-34.16% faster mapping and 1.13-1.26% higher F1-scores for HiFi data (**Figs. 2G and 2H, Table S2**), and 8.14-9.83% faster mapping with 0.92-1.70% higher F1-scores for Cyclone data (**Fig. S5, Table S3**). These findings underscore the adaptability of our Bayesian optimization framework in discovering task-specific, near-optimal parameter configurations that enhance both speed and accuracy.

Furthermore, the utility of our Bayesian optimization framework extends well beyond the specific case of minimap2 alignment. By decoupling the optimization engine from the underlying bioinformatic tool, we establish a tool-agnostic strategy for systematic parameter tuning. This framework is readily adaptable to other computationally intensive workflows, such as de novo assembly or variant calling, contingent only on the definition of a quantifiable objective metric. This represents a shift away from static, "one-size-fits-all" defaults toward dynamic, data-driven pipeline configuration.

## Conclusions

We have developed CycSim, a context-aware simulator that faithfully reproduces the complex error characteristics of long-read sequencing data. We paired this with a analysis-goal-driven Bayesian optimization framework that enables systematic refinement of mapping parameters. Together, these tools provide a robust foundation for improving the accuracy and efficiency of long-read analyses and optimizing bioinformatics workflows for specific platforms and analytical goals.

## Methods

### Model training

Reads shorter than 10 kb were removed using fxTools (v0.3.1, https://github.com/moold/fxTools). The remaining reads from ONT, HiFi, and Cyclone platforms were aligned to the diploid HG002 reference genome using minimap2 (v2.29) with platform-specific presets: ONT (-x lr:hq), HiFi (-x map-hifi), and Cyclone (-k16 -w13 -A2 -B4 -O4,41 -E2,1 -s180 -U70,1000000). Eight Chromosomes (1, 2, 17, and 18 from both haplotypes) and their corresponding reads were extracted for platform-specific model training using different simulators.

For CycSim, ONT and Cyclone reads models were trained using cycsim train -r nanopore -t 30 reads.bam Chr1_2_17_18.fa, while HiFi reads models used cycsim train -r hifi -t 30 reads.bam Chr1_2_17_18.fa. NanoSim (v3.2.3) models were trained with read_analysis.py genome --fastq reads.fastq.gz -rg Chr1_2_17_18.fa -t 30 -c. For Badread (v0.4.1), error and quality score models were generated using badread error_model --reference Chr1_2_17_18.fa --reads reads.fastq.gz --alignment map.paf and badread qscore_model --reference Chr1_2_17_18.fa --reads reads.fastq.gz --alignment map.paf. As PBSIM3 does not provide a training module, the pretrained models supplied by the authors for ONT and HiFi data were used directly.

### Data simulation and evaluation

Simulated reads were generated at 20× depth for the same eight HG002 Chromosomes using each simulator with its corresponding trained model. CycSim reads were produced using cycsim sim -t 30 -d 20 -c model.cy Chr1_2_17_18.fa. NanoSim reads were simulated with simulator.py genome -rg Chr1_2_17_18.fa -c training -x 20 -t 30 --fastq. For Badread, Cyclone reads were generated using badread simulate --reference Chr1_2_17_18.fa --quantity 20x --error_model badread_errors --qscore_model badread_qscore --length 20000,15000 --identity 97,100,2.5. ONT simulations used an adjusted identity range (20, 3), and HiFi simulations used platform-specific length (15374, 13000) and identity (30, 3) settings. For PBSIM3, ONT reads were produced using pbsim --strategy wgs --method errhmm --errhmm ERRHMM-ONT-HQ.model --depth 20 --genome Chr1_2_17_18.fa --length-mean 20000 --accuracy-mean 0.994 --accuracy-min 0.95 --length-min 5000. HiFi reads were generated with pbsim --strategy wgs --method errhmm --errhmm ERRHMM-SEQUEL.model --depth 20 --genome Chr1_2_17_18.fa --length-mean 15374 --pass-num 15, followed by CCS processing (ccs, default parameters) to produce final HiFi reads. All simulated reads were then aligned to the reference genome using minimap2, and we quantified read length, alignment identity, and error biases for each simulator.

### Alignment parameter optimization

CycSim was used to simulate reads with default settings and platform-specific trained models. Simulated ONT, HiFi, and Cyclone datasets were aligned to the reference genome using minimap2. SVs were called using Sniffles2 [13] (v2.6.3) with the --tandem-repeats option, and benchmarked against the HG002 GIAB v1.1 truth set [11] using truvari [14] with --passonly -r 1000 -p 0.00 --refine. SNP calling was performed with Longshot [15], and evaluated using hap.py with the --engine vcfeval (https://github.com/Illumina/hap.py).

The optimal minimap2 parameters identified for SV calling using ONT data were -k 21 -w 21 -A 1 -B 9 -O 13,44 -E 3,1 -s 30 -U 20,50000. For HiFi data SV calling, the best parameters were -k 23 -w 22 -A 1 -B 9 -O 13,41 -E 3,1 -s 180 -U 10,5000. For Cyclone data, the optimal parameters for general alignment were -w 13 -A 2 -B 4 -O 4,41 -E 2,1 -s 180 -U70,1000000, whereas the optimal parameters specifically for SV calling were -k 17 -w 13 -A 1 -B 9 -O 13,44 -E 3,1 -s 30 -U70,500.

## Declarations

### Ethics approval and consent to participate

Not applicable.

### Consent for publication

Not applicable.

### Availability of data and materials

The HG002 v1.1 reference genome and HiFi sequencing data were obtained from the HG002 repository (https://github.com/marbl/HG002).

The HG002 ONT Q20 dataset was downloaded from the ONT Open Datasets portal (ont-open-data.s3.amazonaws.com/gm24385_2023.12/all_pass.vhg002v1. bam). HG002 Cyclone reads were retrieved from the China National GeneBank (CNGB) under accession CNP0007646. CycSim and its pretrained models are released under the Massachusetts Institute of Technology (MIT) License and are available on GitHub [16] and Zenodo [17].

### Authors’ contributions

JH and CY jointly designed and supervised the project. JH developed CycSim and drafted the manuscript. DF and XJ review the code and provided the suggestions. All authors contributed to manuscript revision and approved the final version.

## Acknowledgements

We thank Tao Zeng, Huiyuan Hao, and Jiayuan Zhang for providing the HG002 Cyclone data.

## Supplementary Information

**Figure S1.**
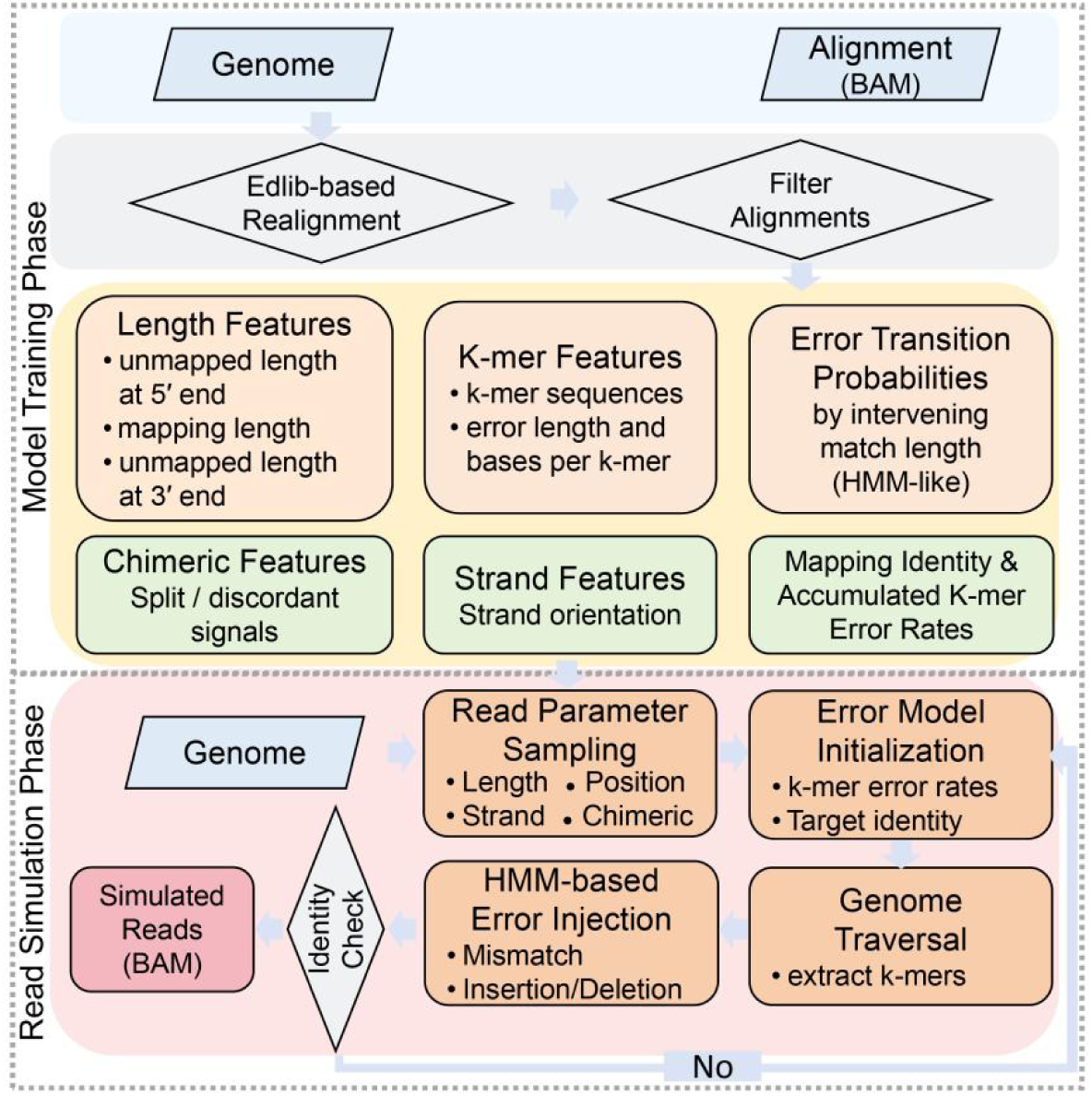
CycSim pipeline. The CycSim pipeline operates in two stages: model training and read simulation. The training stage processes high-confidence BAM alignments to characterize read structure and models error characteristics at two complementary levels: K-mer-based modeling and Error transition modeling. In the simulation stage, these derived models define the read’s structure and error profile, and the aligned read core is generated via a base-wise sliding process that samples errors using both models. A detailed description is available in the "Context-aware long-read simulation" section.

**Figure S2.**
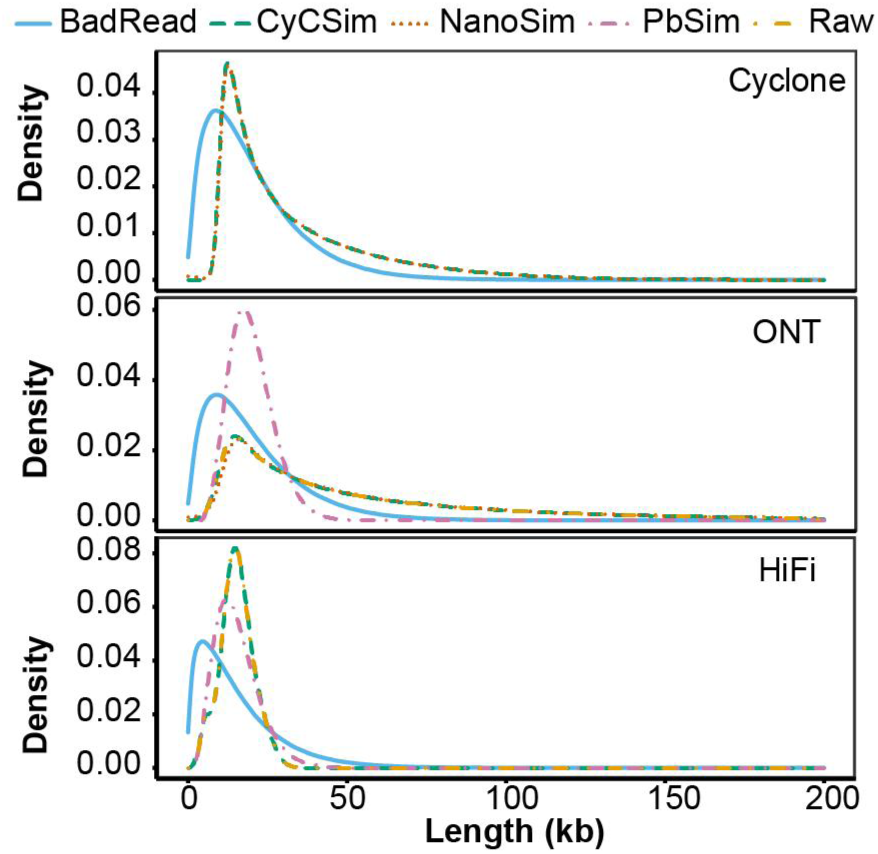
Distribution of reads length of simulated reads compared with real reads.

**Figure S3.**
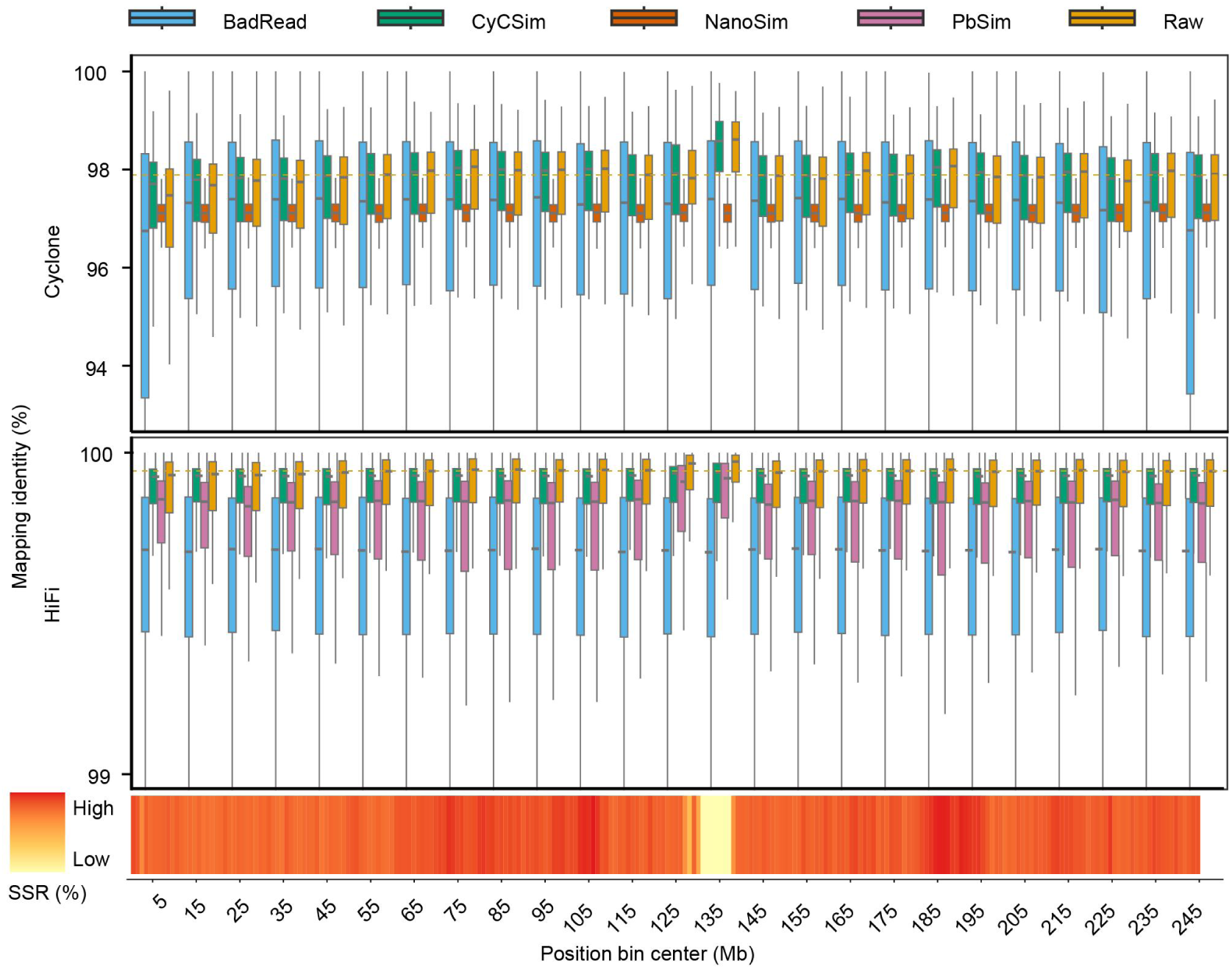
Positional alignment identity distribution along Chr1 maternal for simulated Cyclone and HiFi reads. Horizontal yellow lines mark the median identity of real ONT reads. The lowest heatmap shows the short tandem repeats (STRs) density (1–6 bp motifs, ≥3 repeat units) along Chr1 maternal.

**Figure S4.**
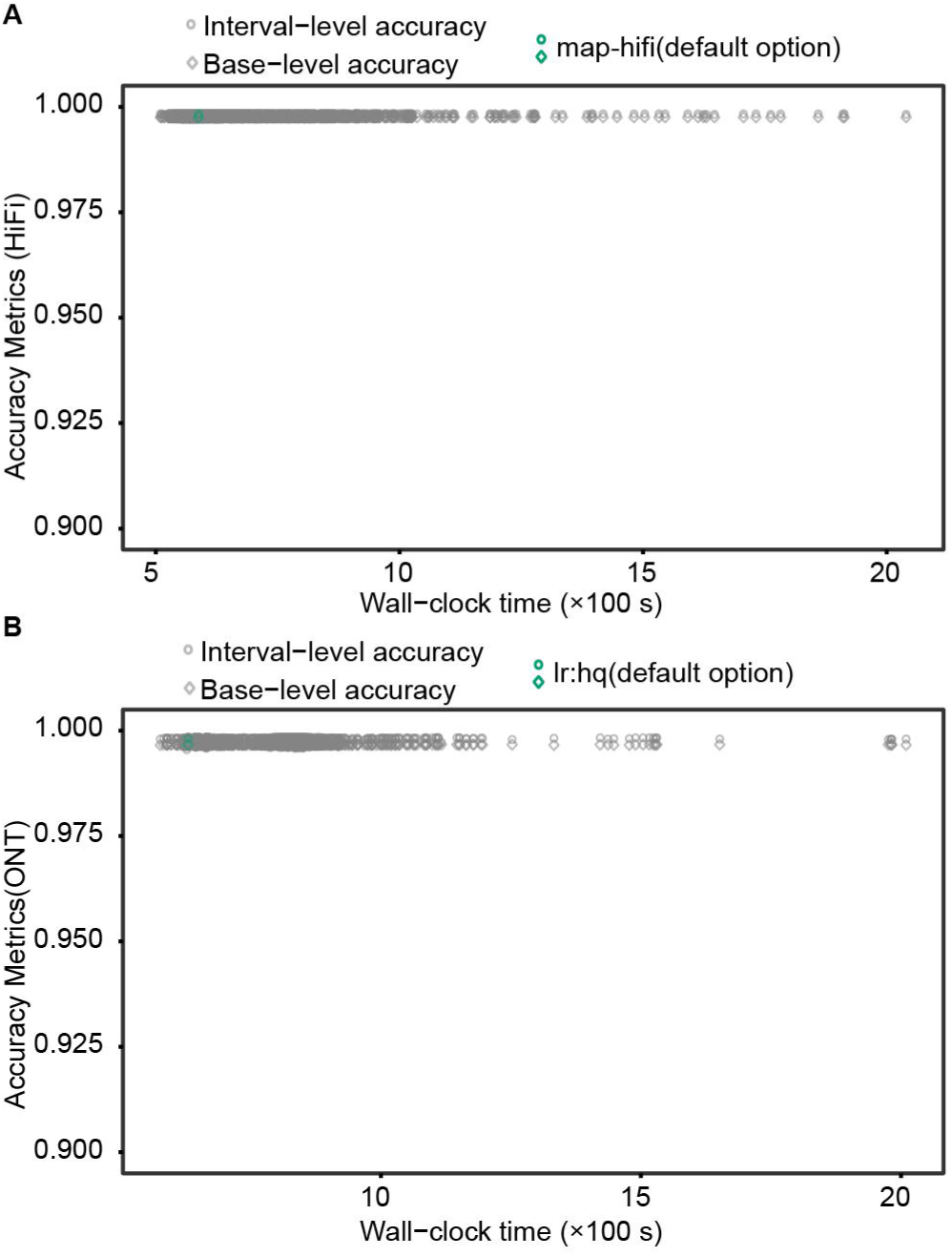
Performance of HiFi and ONT mapping parameters on simulated reads. Interval-level accuracy denotes the proportion of aligned intervals within 50 bp of the true interval, and Base-level accuracy denotes the proportion of correctly aligned bases.

**Figure S5.**
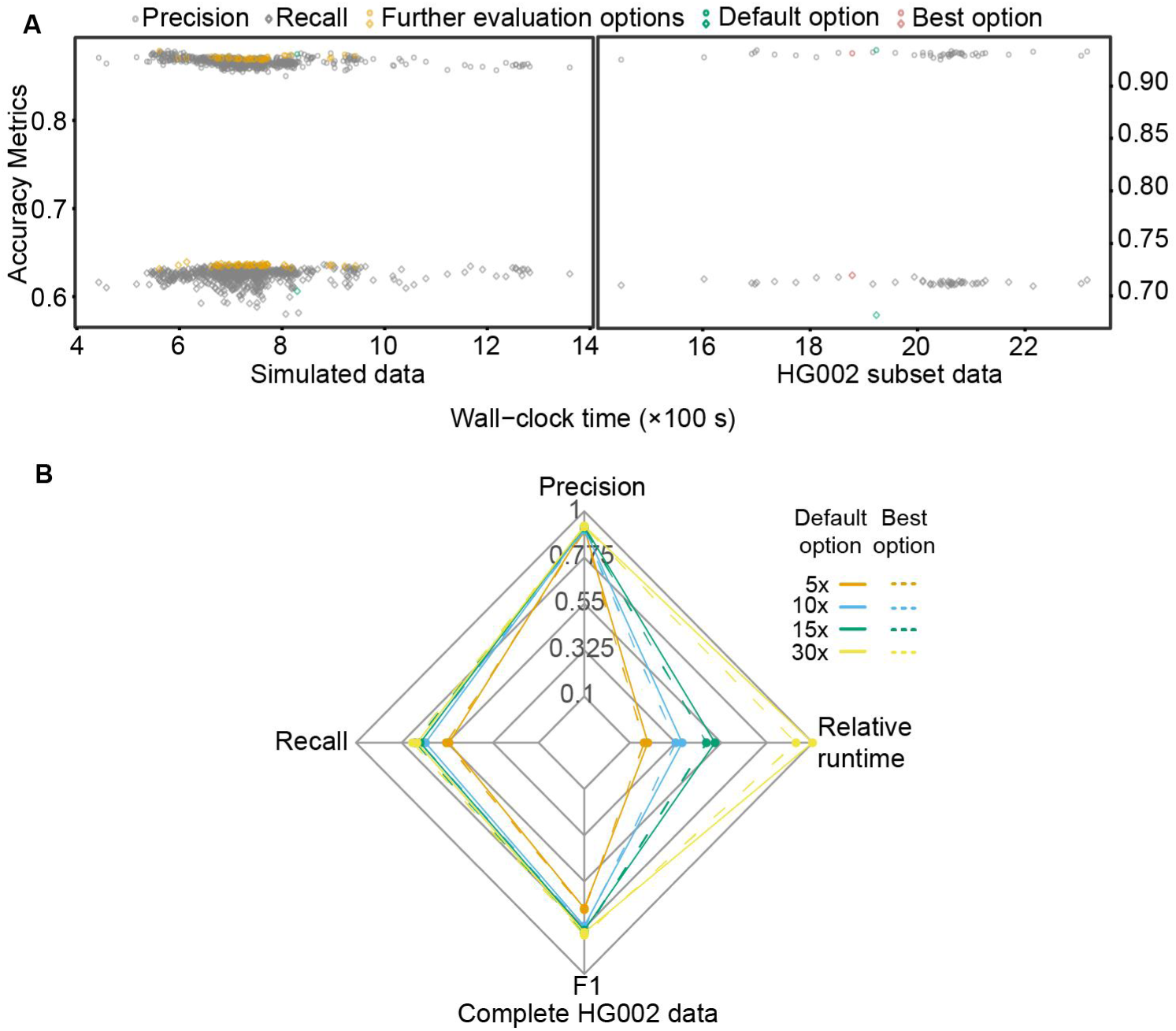
Bayesian optimization and evaluation of mapping parameters. **A** Performance of Cyclone SV detection-oriented mapping parameters on simulated and partial real reads. Parameters evaluated in the right panel were selected from the left panel (labeled as Further evaluation options), and Wall-clock time includes both minimap2 and samtools sort. The default option represents the optimal parameters we identified for general-purpose mapping. **B** Performance of optimized versus default Cyclone mapping parameters for SV detection at different coverage depths. Relative runtime denotes the proportion of runtime relative to the longest run, considering minimap2 only.

**Table S1.** Performance of optimized versus default ONT mapping parameters for SV detection at different coverage depths.

| Depth | Options | Precision | Recall | F1 score | Wall-clock time | Relative runtime |
| --- | --- | --- | --- | --- | --- | --- |
| 5x | Best option | 0.9199 | 0.5817 | 0.7127 | 480.6460 | 0.1369 |
| 5x | lr:hq (default) | 0.9360 | 0.5653 | 0.7049 | 695.0120 | 0.1979 |
| 10x | Best option | 0.9159 | 0.7076 | 0.7984 | 851.2320 | 0.2424 |
| 10x | lr:hq (default) | 0.9341 | 0.6881 | 0.7924 | 1263.9450 | 0.3599 |
| 15x | Best option | 0.9276 | 0.7373 | 0.8216 | 1275.1790 | 0.3631 |
| 15x | lr:hq (default) | 0.9369 | 0.7196 | 0.8140 | 1853.2520 | 0.5277 |
| 30x | Best option | 0.9395 | 0.7553 | 0.8374 | 2369.3420 | 0.6747 |
| 30x | lr:hq (default) | 0.9390 | 0.7374 | 0.8261 | 3511.8560 | 1.0000 |

**Table S2.**
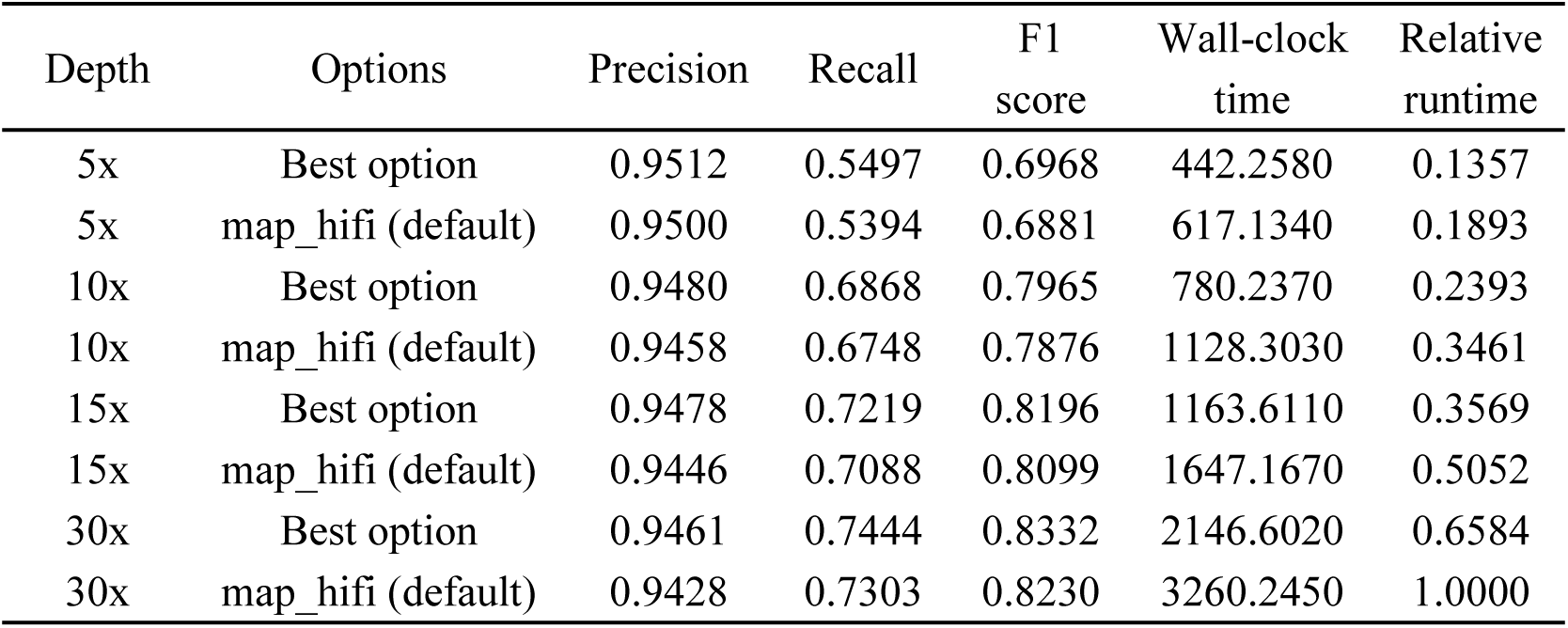
Performance of optimized versus default HiFi mapping parameters for SV detection at different coverage depths.

**Table S3.** Performance of optimized versus default Cyclone mapping parameters for SV detection at different coverage depths.

| Depth | Options | Precision | Recall | F1 score | Wall-clock time | Relative runtime |
| --- | --- | --- | --- | --- | --- | --- |
| 5x | Best option | 0.9009 | 0.5548 | 0.6867 | 638.6830 | 0.1698 |
| 5x | default option | 0.9157 | 0.5414 | 0.6804 | 708.3230 | 0.1883 |
| 10x | Best option | 0.9057 | 0.6820 | 0.7781 | 1226.8020 | 0.3261 |
| 10x | default option | 0.9180 | 0.6578 | 0.7664 | 1344.6330 | 0.3574 |
| 15x | Best option | 0.9197 | 0.7083 | 0.8002 | 1793.3320 | 0.4767 |
| 15x | default option | 0.9266 | 0.6838 | 0.7869 | 1952.3460 | 0.5190 |
| 30x | Best option | 0.9249 | 0.7217 | 0.8108 | 3454.6210 | 0.9183 |
| 30x | default option | 0.9282 | 0.6993 | 0.7977 | 3761.9850 | 1.0000 |

The default option represents the optimal parameters we identified for general-purpose mapping.

